# Porcine placenta extract upregulates ceramide synthase 3 expression via the PPARδ/ILK/Akt/mTOR/STAT3 pathway

**DOI:** 10.1101/2022.05.23.493164

**Authors:** Akihiro Aioi, Ryuta Muromoto, Jun-ichi Kashiwakura, Sho Mogami, Megumi Nishikawa, Shigeyuki Ogawa, Tadashi Matsuda

## Abstract

Porcine placenta extract (PPE) is widely accepted as an ingredient in complementary and alternative medicine and previous studies have reported its availability, however, its underlying action mechanism remains unclear. In this study, we investigated the underlying mechanism of porcine placenta extract-induced ceramide synthase 3 upregulation. PPE enhanced the expression of ceramide synthase 3 at both the mRNA and protein levels in HaCaT cells. Moreover, porcine placenta extract-induced ceramide synthase3 upregulation was suppressed by the Akt inhibitor, suggesting the involvement of Akt in the underlying mechanism. As the PI3K inhibitor did not affect porcine placenta extract-induced ceramide synthase 3 upregulation, the factors upstream of Akt were estimated. Inhibition and small interfering RNA experiments demonstrated that the peroxisome proliferator-activated receptors δ (PPAR δ) and integrin-linked kinase (ILK) are involved in the phosphorylation of Akt. Next, we explored the factors downstream of Akt and found that porcine placenta extract induced phosphorylation of the STAT3 while porcine placenta extract-induced upregulation of ceramide synthase 3 was significantly suppressed by the inhibitor of the mTOR, suggesting that the mTOR/STAT3 pathway is involved in the downstream of Akt. These results demonstrate that porcine placenta extract upregulates CerS3 expression via the PPARδ/ILK/Akt/mTOR/STAT3 pathway.

## Background

The primary function of the epidermis is to construct an indispensable barrier between the organism and the desiccated terrestrial environment preventing invasion of external stimuli and loss of water from the body. As summarized in a previous study, many factors are involved in epidermal barrier function [1, 2]. Lipids, which are stored in lamellar bodies in the stratum granulosum and secreted into the extracellular space, are crucial for epidermal barrier function [3]. Previous studies have reported that lipid synthesis is regulated by various inter- and intracellular signaling pathways. Epidermal barrier disruption induces acute upregulation of the expression of inflammatory cytokines in the epidermis, although a chronic increase in cytokine production could have a harmful effect leading to inflammation and epidermal hyperproliferation [4]. Previous studies have suggested that inflammatory cytokines such as interleukin (IL)-6 and tumor necrosis factor (TNF) are potent stimulators of lipid synthesis to maintain epidermal homeostasis [5, 6]. In contrast, previous studies have demonstrated that peroxisome proliferator-activated receptor (PPAR) activators not only significantly enhance the synthesis of lipids including barrier-specific ceramide species, but also significantly accelerate the lamellar body formation, secretion, and post-secretory processing [7]. In addition, Mizutani *et al*. reported that activators of PPARβ/δ and PPARγ induce ceramide synthase 3 (CerS3) and that PPARβ/δ small interfering RNA (siRNA) inhibits the induction of CerS3 mRNA during differentiation [8]. These results suggest that PPARs play roles in the maintenance of epidermal barrier homeostasis.

Clinical applications of human and/or porcine placenta extract (PPE) are widely accepted in oriental medicine because the placenta is a rich reservoir of diverse bioactive molecules. A previous study showed that injection of human placenta extract into wound boundaries accelerated the decrease in wound size [9]. Moreover, oral administration of PPE reportedly reduces UVB-induced wrinkle formation in mice, along with the downregulation of matrix metalloproteinase (MMP)-2 mRNA expression [10]. In addition, studies using skin cells have revealed that human placenta extract accelerates keratinocyte and fibroblast proliferation [11, 12]. Recently, we reported that the application of PPE-containing gel on the face alleviated wrinkle formation and improved skin hydration, along with the upregulation of CerS3 in keratinocytes [13]. However, as the underlying mechanism still remains unclear, we explored the signaling pathway of PPE-induced CerS3 induction in this study.

## Materials and Methods

### Preparation of PPE

PPE was kindly provided by Kasyu Industries (Kitakyushu, Fukuoka, Japan). Briefly, fresh porcine placenta, collected at calving at the farm, was washed with water and then subjected to the freeze (−20°C)-thawing (under 10°C) cycle. After pasteurization at 65°C for 6 h and removal of debris with 0.22δm filter, the obtained freeze-thaw drip was diluted with sterilized water to a nitrogen content of 0.27% to 0.33%, according to the Kjeldahl method described in the Japanese Standards of Food Additive.

### Reagents and antibodies

All inhibitors and an activator were purchased from Selleck Chemicals (Houston, TX, USA). Primary antibodies against β-actin antibody, CerS3, p-Akt (Ser473) and p-STAT3 were purchased from Santa Cruz Biotechnology (Santa Cruz, CA, USA), Thermo Fisher Scientific (Waltham, MA, USA), and Cell Signaling Technology (Beverly, MA, USA), respectively.

### Cells

To maintain the differentiation stage of HaCaT cells, calcium in fetal bovine serum was depleted by incubation with Chelex 100 resin (BioRad, Hercules, CA, USA) for 1 h at 4°C. HaCaT cells were maintained in Ca^2+^-free Dulbecco’s modified Eagle’s medium (DMEM) supplemented with 5% Ca^2+^-depleted fetal bovine serum, 4 mM glutamine, 1 mM sodium pyruvate and 2 mM CaCl_2_ at 37°C in a 5% carbon dioxide (CO_2_-)-humidified atmosphere.

### Treatment with PPE and reagents

HaCaT cells were seeded into 24-well plates at a cell density of 1 × 10^5^ cells/well for qPCR and into 6-well plates at a cell density of 3 × 10^5^ cells/well for immunoblotting, and then maintained in a 5% CO_2_-humidified atmosphere at 37°C. After cultivation for 24 h, the cells were treated with 5% PPE in the presence or absence of reagents for appropriate periods.

### siRNA transfection

HaCaT cells were reverse-transfected with predesigned PPARδ, and ILK siRNA (Thermo Fisher Scientific, Waltham, MA), according to the manufacturer’s instructions. Cells were seeded into 24-well plates at a cell density of 1 × 10^5^ cells/well for qPCR and into 6-well plates at a cell density of 3 × 10^5^ cells/well for immunoblotting, and incubated in a humidified atmosphere of 5% CO_2_ at 37°C for 24 h, followed by treatment with PPE and reagents for appropriate periods. The harvested cells were subjected to qPCR and immunoblotting assay.

### RNA isolation and qPCR

After the treatments, cells were harvested and total RNA was prepared with SV RNA isolation kit (Promega, Madison, WI, USA), according to the manufacturer’s instructions, followed by reverse transcription using ReverTra Ace® qPCR RT Master Mix (TOYOBO, Osaka, Japan). PCR amplification and detection were conducted on a CFX96 real-time PCR system (BioRad, Hercules, CA, USA) using a KAPA SYBR FAST qPCR master mix (KAPA Biosystems, Woburn, MA, USA). The following primer pairs were used: β-actin, 5’-GATGAGATTGGCATGGCTTT-3’ (sense) and 5’-CACCTTCACCGTTCCAGTTT-3’ (antisense); CerS3, 5’- ACATTCCACAAGGCAACCATTG-3’ (sense) and 5’-CTCTTGATTCCGCCGACTCC- 3’ (antisense); PPARδ, 5’- ACTGAGTTCGCCAAGAGCAT-3’ (sense) and 5’- TGCACGCCATACTTGAGAAG-3’ (antisense); ILK, 5’- AAGGTGCTGAAGGTTCGAGA-3’ (sense) and 5’- ATACGGCATCCAGTGTGTGA- 3’ (antisense). The expression of target mRNA was quantified using the comparative threshold cycle (Ct) method for relative quantification (2^-δδCt^), normalized to the geometric mean of the reference gene β-actin.

### Immunoblotting

HaCaT cells were collected in the RIPA buffer supplemented with protease inhibitors and phosphatase inhibitors. Equal amount of protein (10μg) were loaded, resolved via SDS-PAGE and transferred to PVDF membrane, followed by immunoblotting with each antibody. Immunoreactive proteins were visualized using an enhanced chemiluminescence detection system (Millipore, Bedford, MA, USA). The relative intensity of the blot was measured using the ImageJ software and was normalized by the intensity of β-actin blot.

### Statistical analysis

Results of mRNA relative expression and densitometric analysis of western blotting images are expressed as the mean ± standard deviation (SD) of at least three independent experiments. Statistical analysis was performed using the Student’s t-test. Statistical significance was set *p* < 0.05.

## Results

### PPE enhanced CerS3 expression

To confirm the effect of PPE on the mRNA expression of CerS3, quantitative polymerase chain reaction (qPCR) was performed. The treatment with PPE significantly upregulated the mRNA expression of CerS3 to 2.40 ± 0.59 (Figure 1a). As well, the protein expression of CerS3 was significantly enhanced to 2.00 ± 0.58 (Figure 1b and 1c).

**Figure 1.**
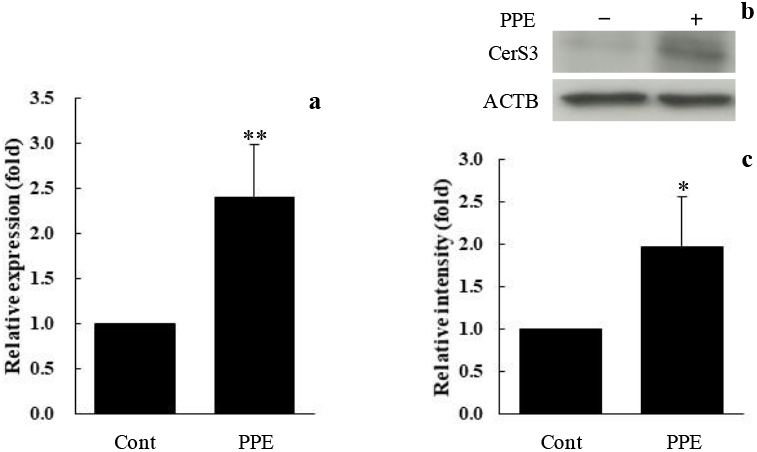
Porcine placenta extract (PPE) enhanced ceramide synthase 3 (CerS3) expression. (a) CerS3 mRNA expression was upregulated in PPE-treated cells. (b) Representative western blotting image of equal amounts of protein extracts from the control and PPE-treated cells. (c) Relative intensity of CerS3 blots was measured using the ImageJ software and was normalized by the intensity of the β-actin blot. Statistical analyses were performed with a Student’s t-test. * p-value < 0.05, ** p-value < 0.01 were considered to be significantly different from the control.

### Akt was involved in PPE-induced upregulation of CerS3

GSK690693, an inhibitor of Akt, was employed to evaluate the involvement of Akt in PPE-enhanced CerS3 mRNA and protein expression. Treatment with GSK690693 significantly downregulated PPE-enhanced CerS3 mRNA expression (Figure 2a). In addition, the protein level of CerS3 was significantly decreased by treatment with GSK690693 (Figure 2b and 2c). In contrast, LY294002, an inhibitor of phosphoinositide 3-kinase (PI3K) that phosphorylates Akt, did not affect PPE-induced upregulation of CerS3 expression (Figure 2d).

**Figure 2.**
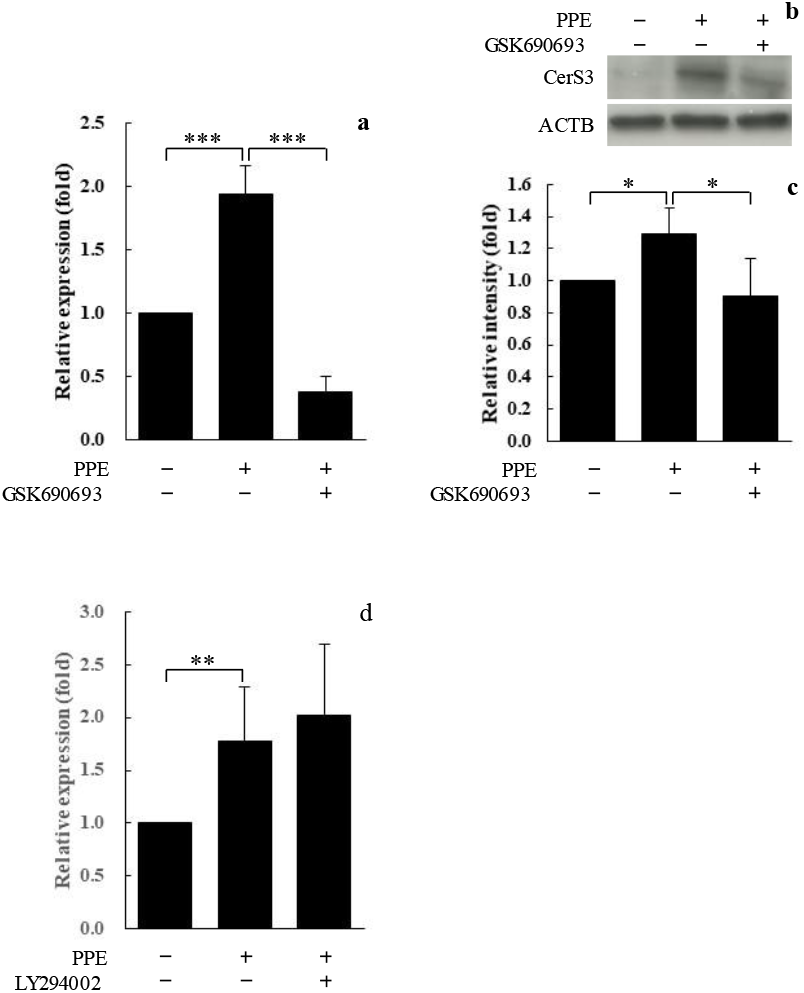
Akt was involved in PPE-induced upregulation of CerS3 expression, but not PI3K. (a) GSK690693 suppressed CerS3 mRNA expression upregulated by PPE treatment. (b) Representative western blotting image of equal amounts of protein extracts from the control and PPE-treated cells in the absence and presence of GSK690693. (c) Relative intensity of CerS3 blots was measured using the ImageJ software and was normalized by the intensity of the β-actin blot. (d) LY294002 did not affect PPE-upregulated CerS3 mRNA expression. Statistical analyses were performed with a Student’s t-test. * p-value < 0.05, ** p-value < 0.01 and *** p-value < 0.001were considered to be significantly different from the control.

### PPARδ was involved in PPE-induced upregulation of CerS3

The PPARδ selective inhibitor, GSK3787, significantly downregulated PPE-enhanced CerS3 mRNA expression, whereas GSK9662, a selective inhibitor of PPARδ, did not affect PPE-enhanced CerS3 mRNA expression (Figure 3a). Moreover, GW0742, a PPARδ activator, upregulated CerS3 mRNA expression as well as PPE (Figure 3b). While the expression of PPARδ was knocked down by siRNA (Figure 3c), the effect of PPE on CerS3 expression was diminished (Figure 3d). In addition, GSK3787 and PPARδ knockdown (KD) significantly reduced the upregulation of CerS3 protein levels by PPE treatment (Figure 3e-3h).

**Figure 3.**
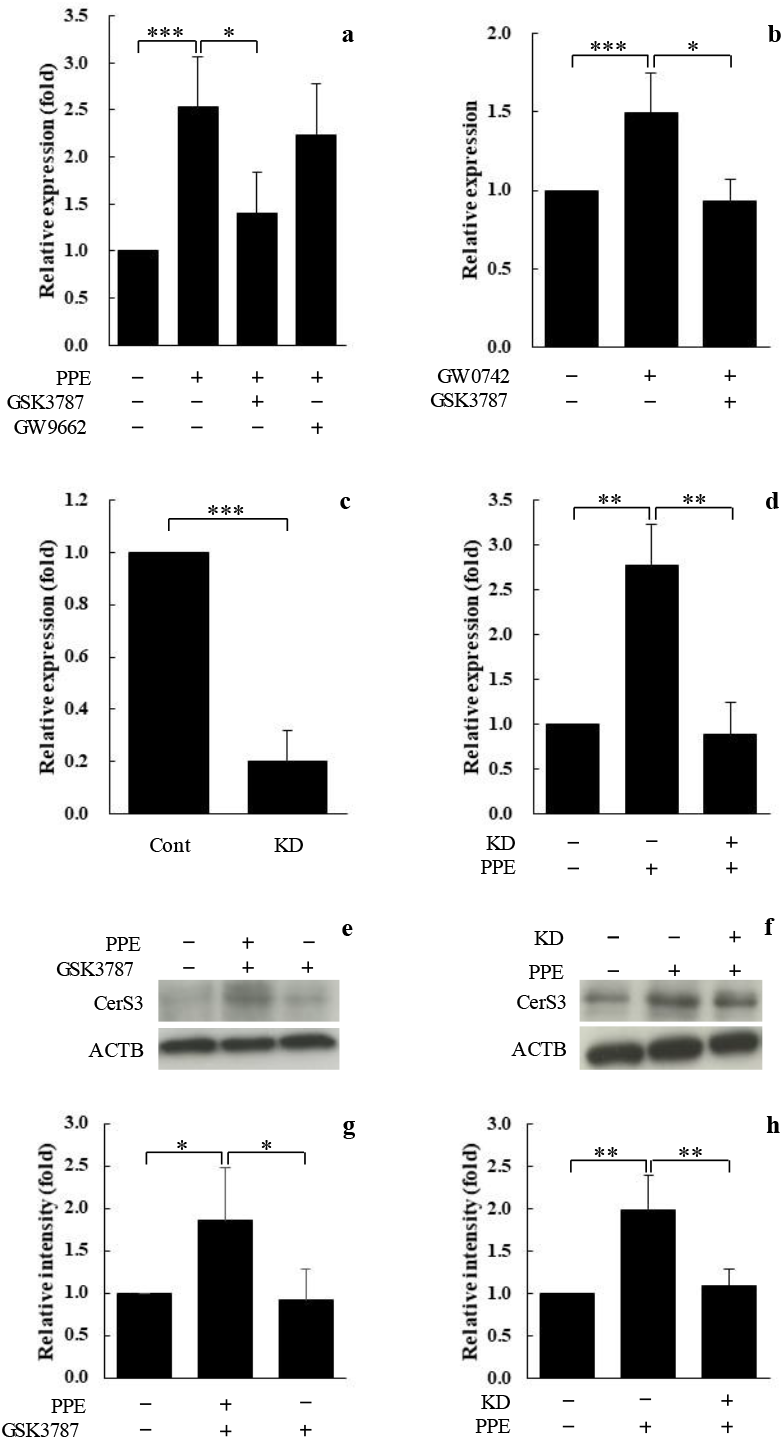
PPARδ was involved in PPE-induced upregulation of CerS3 expression, but not PPARδ. (a) GSK3787 suppressed CerS3 mRNA expression upregulated by PPE treatment, while GW9662 did not affect the upregulation. (b) GW0742 enhanced CerS3 mRNA expression and its upregulation was suppressed by GSK3787. (c) Expression of PPARδ mRNA was reduced by the small interfering RNA (siRNA). (d) PPARδ knockdown (KD) suppressed CerS3 mRNA expression upregulated by PPE treatment. (e) Representative western blotting image of equal amounts of protein extracts from the control and PPE-treated cells in the absence and presence of GSK3787. (f) Representative western blotting image of equal amounts of protein extracts from the control and PPARδ KD cells. (g and h) Relative intensity of CerS3 blots was measured using the ImageJ software and was normalized by the intensity of the β-actin blot in inhibition and siRNA experiments. Statistical analyses were performed with a Student’s t-test. * p-value < 0.05, ** p-value < 0.01 and *** p-value < 0.001were considered to be significantly different from the control.

### Integrin-kinked kinase (ILK) was involved in PPE-induced upregulation of CerS3

The ILK inhibitor, OSU-T315, drastically reduced the effect of PPE on CerS3 expression at both the mRNA and protein levels (Figure 4a-4c). While the expression of ILK was knocked down by siRNA (Figure 4d), the upregulation of CerS3 expression by PPE was diminished significantly at both the mRNA and protein levels (Figure 4e-4g). The Mrna expression of ILK was significantly enhanced after 16 h of PPE treatment, following significant upregulation of PPARδ mRNA expression (Figure 4h).

**Figure 4.**
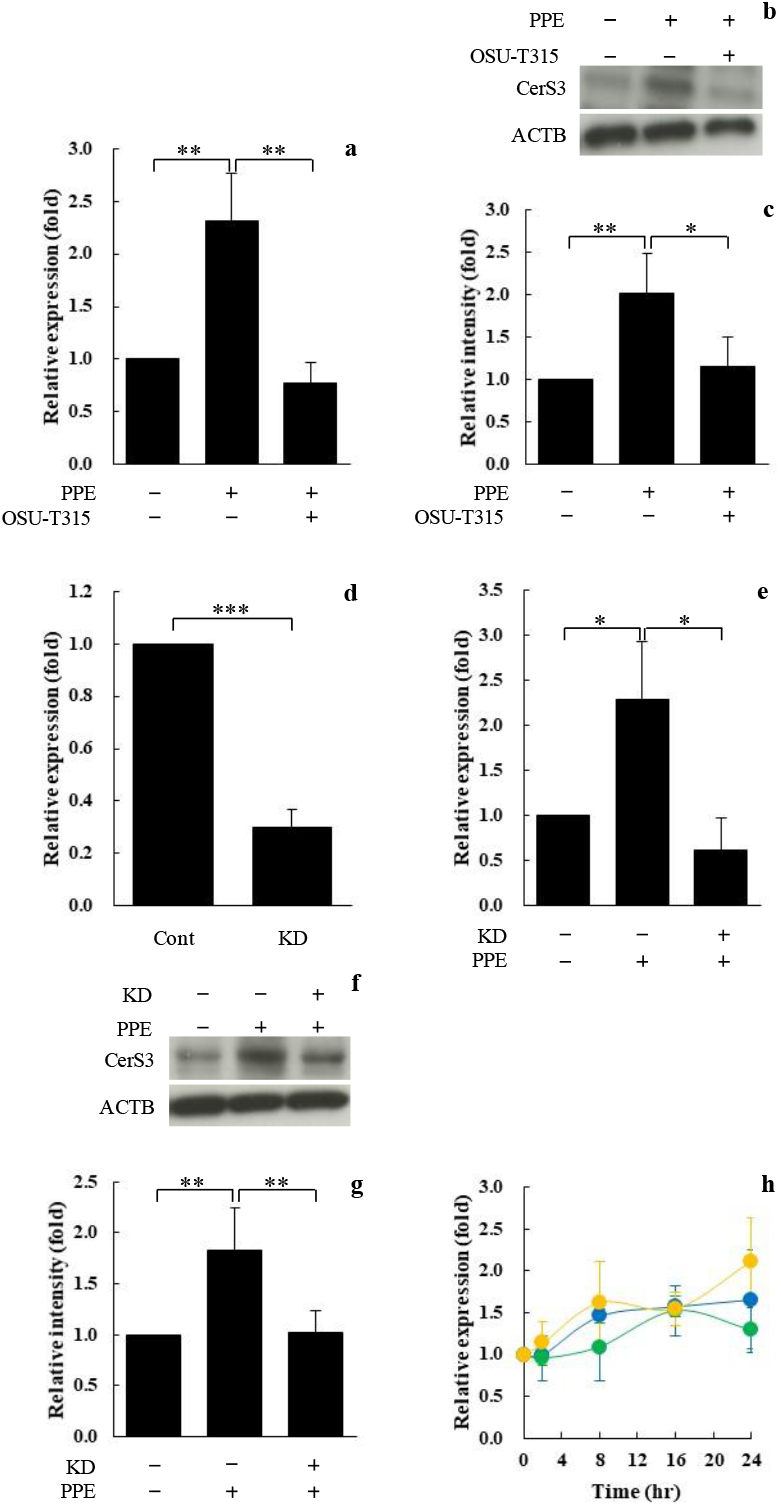
ILK was involved in PPE-induced upregulation of CerS3 expression. (a) OSU-T315 suppressed CerS3 mRNA expression upregulated by PPE treatment. (b) Representative western blotting image of equal amounts of protein extracts from the control and PPE-treated cells in the absence and presence of OSU-T315. (c) Relative intensity of CerS3 blots was measured using the ImageJ software and was normalized by the intensity of the β-actin blot in the inhibition experiments. (d) Expression of ILK mRNA was reduced by siRNA. (e) ILK KD suppressed CerS3 mRNA expression upregulated by PPE treatment. (f) Representative western blotting image of equal amounts of protein extracts from the control and the ILK KD cells. (g) Relative intensity of CerS3 blots was measured using the ImageJ software and was normalized by the intensity of the β-actin blot in the siRNA experiments. (h) CerS3 mRNA expression (yellow circle) arose in the acute phase (after 2hr) and was maintained throughout the experimental period. PPARδ mRNA expression (blue circle) arose after 8hr, followed by ILK mRNA expression (green circle). Statistical analyses were performed with a Student’s t-test. * p-value < 0.05, ** p-value < 0.01 and *** p-value < 0.001were considered to be significantly different from the control.

### mTOR/STAT3 signaling pathway participated in PPE-induced upregulation of CerS3

The PPE-induced CerS3 expression at the mRNA and protein levels was significantly reduced in the presence of rapamycin, an mTOR inhibitor (Figure 5a-5c). Phosphorylated STAT3 (p-STAT3) was detected in HaCaT cells treated with PPE, as well as phosphorylated Akt (Figure 5d), suggesting the involvement of STAT3 in PPE-induced CerS3 upregulation.

**Figure 5.**
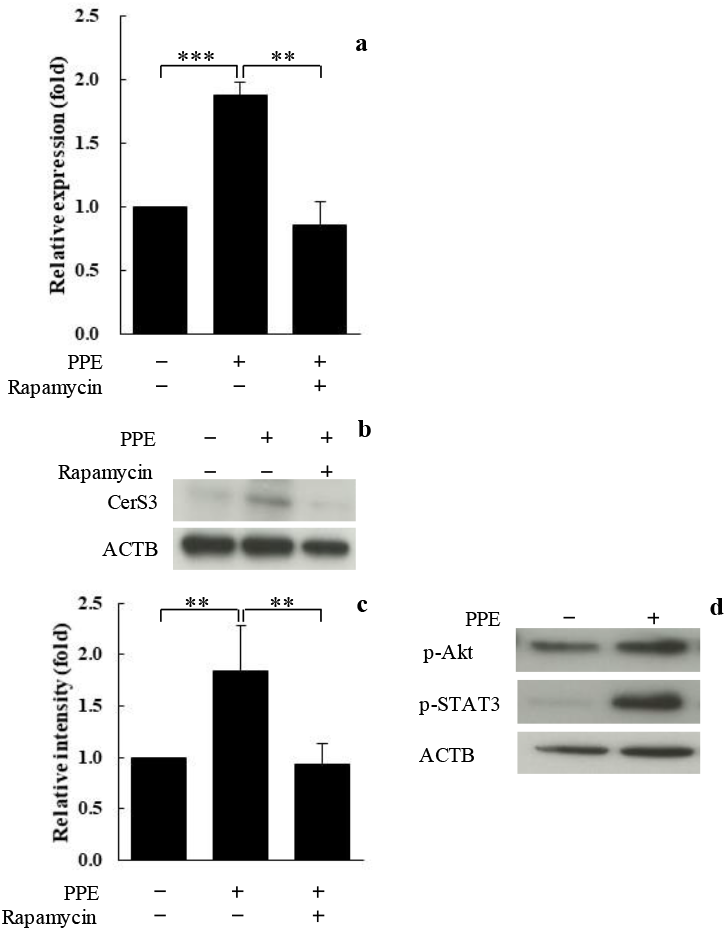
mTOR)/STAT3 signaling pathway was involved in PPE-induced upregulation of CerS3 expression. (a) Rapamycin, an mTOR inhibitor, suppressed CerS3 mRNA expression upregulated by PPE treatment. (b) Representative western blotting image of equal amounts of protein extracts from the control and PPE-treated cells in the absence and presence of rapamycin. (c) Relative intensity of CerS3 blots was measured using the ImageJ software and was normalized by the intensity of the β-actin blot in inhibition experiments. (d) Representative western blotting image of p-Akt and p-STAT3 in the cells treated with PPE for 30 min. Statistical analyses were performed with a Student’s t-test. * p-value < 0.05, ** p-value < 0.01 and *** p-value < 0.001were considered to be significantly different from the control.

## Discussion

Ceramides, which are the predominant lipids comprising 30 to 50 % of the intercellular lipid content in the stratum corneum, are synthesized by the enzymatic actions of ceramide synthases (CerS). CerS3, which is one of the six distinct isoforms of CerSs, is highly expressed in keratinocytes, synthesizes ceramides containing α-hydroxy fatty acids, and is involved in maintaining skin barrier function [14-16]. We have recently reported that PPE, which is widely used in alternative and traditional medicine, enhances CerS3 expression [13]. However, because the regulation of CerS3 expression is largely uncharacterized, we explored the underlying mechanism of the PPE-induced upregulation of CerS3 in this study. The expression levels of CerS3 mRNA and protein were significantly upregulated in the cells treated with PPE (Figure 1). The upregulation was diminished by Akt inhibitor (Figure 2a and 2b), although LY294002, a PI3K inhibitor, did not affect the effect of PPE (Figure 2c). This suggests that some signaling pathways other than PI3K/Akt are involved in the mechanism underlying the induction of CerS3. To elucidate the upstream and downstream pathways of Akt, we first examined the involvement of PPARδ in the signaling pathway. As shown in Figure 3, PPARδ is involved in the PPE-induced upregulation of CerS3 expression. This corresponds to a previous study demonstrating that PPARβ/δ is involved in CerS3 upregulation during keratinocyte differentiation [8]. However, to the best of our knowledge, the binding site of PPARδ does not exist in the promoter region of the *CerS3* gene, suggesting that PPARδ contributes indirectly to CerS3 upregulation. As described above, Akt is crucial in the signaling pathway involved in PPE-induced CerS3 upregulation, as well as PPARδ. A previous study demonstrated that ILK, which is ubiquitously expressed, phosphorylates Akt at serine 473, leading to the activation of Akt [17, 18]. The inhibition of ILK and ILK KD diminished PPE-induced CerS3 upregulation (Figure 4a-g), suggesting that ILK also plays a pivotal role in the signaling pathway of the effect of PPE. Di-Poï *et al*. reported that PPARβ/δ modulates Akt activation via the upregulation of ILK [19]. As shown in Figure 4h, PPE treatment enhanced PPARδ mRNA expression, followed by the upregulation of ILK mRNA expression, suggesting that enhancement of the expression of PPARδ and ILK contributes to maintaining the upregulated level of CerS3 expression during the experimental period. Collectively, the PPE treatment regulates Akt phosphorylation via ILK whose expression is enhanced by PPARδ upstream of the signaling pathway of PPE-induced CerS3 upregulation. An important effector of Akt via tuberous sclerosis (TSC)-1/2 and the small GTPase Rheb is the mTOR signaling pathway [20, 21]. Therefore, we elucidated the involvement of mTOR in the signaling pathway involved in PPE-induced CerS3 upregulation. Rapamycin, an mTOR inhibitor, suppressed PPE-upregulated CerS3 expression at both the mRNA and protein levels (Figure 5a-c), suggesting the involvement of mTOR in the signaling pathway. STAT3 is a crucial regulator of various cell processes. STAT3 is selectively activated by stimulation with cytokines and growth factors [22, 23]. In keratinocytes, STAT3 plays a crucial role in keratinocyte migration and skin remodeling [24]. mTOR also positively regulates STAT3 activation, similar to cytokines and growth factors [25]. Collectively, the mTOR/STAT3 signaling pathway participates downstream of Akt phosphorylation in PPE-upregulated CerS3 expression. In conclusion, we found that PPE upregulated CerS3 expression via the PPARδ/ILK/Akt/mTOR/STAT3 signaling pathway (Figure 6). In particular, PPE induced the persistent expression of PPARδ, while ILK contributed to the enhanced expression of CerS3.

**Figure 6.**
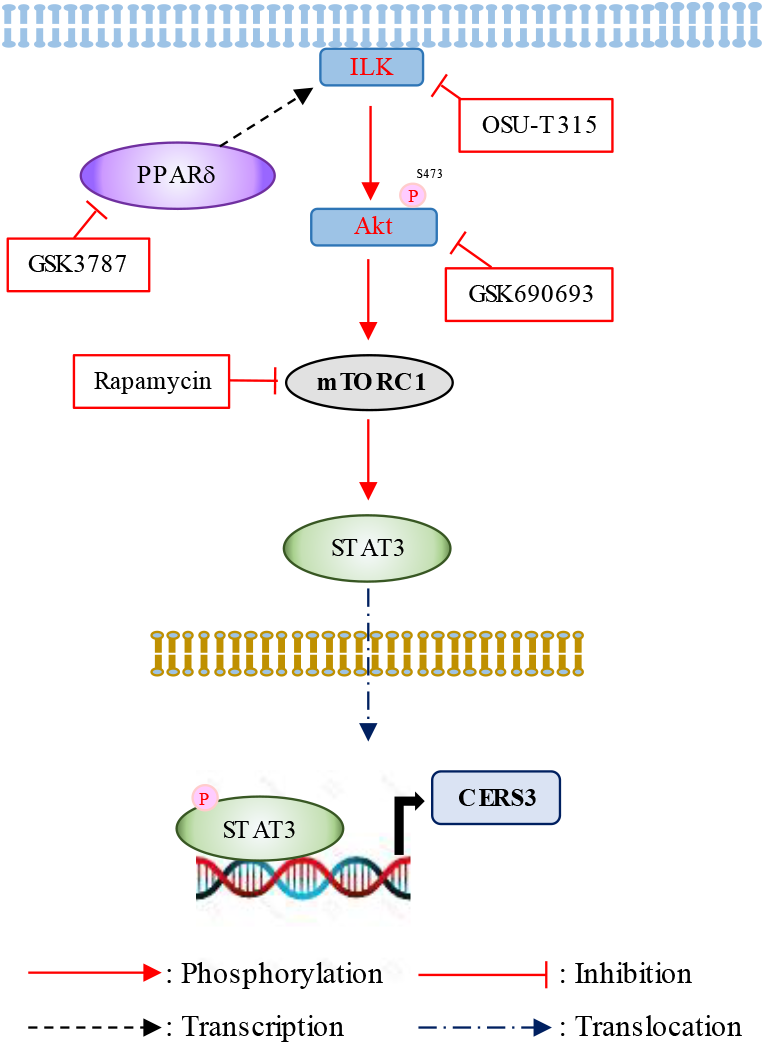
Potential signaling pathway underlying the PPE-induced upregulation of CerS3.

## Conclusion

In this study, we demonstrated, for the first time, the underlying mechanism of PPE function and provided scientific evidence for the application of PPE in alternative medicine.

## List of abbreviations

Cers3: ceramide synthase 3
DMEM: Dulbecco’s modified Eagle’s medium
IL-6: interleukin-6
ILK: integrin-linked kinase
MMP-2: matrix metalloproteinase-2
mTOR: mammalian target of rapamycin
PI3K: phosphoinositide 3-kinase
PPAR: peroxisome proliferator-activated receptor
PPE: porcine placenta extract
PVDF: poly vinylidene difluoride
qPCR: quantitative polymerase chain reaction
RIPA: radio-immunoprecipitation assay
siRNA: small interfering RNA
SDS-PAGE: sodium dodecyl sulfate-polyacrylamide gel electrophoresis
STAT3: signal transducer and activator of transcription 3
TNF: tumor necrosis factor

## Declarations

### Availability of data and materials

All data generated or analyzed during this study are included in this published article.

## Acknowledgements

We would like to thank Editage (www.editage.com) for English language editing

## References

1 Natsuga K. Epidermal barriers. Cold Spring Harb Perspect Med. 2014; doi: 10.1101/cshperspect.a018218

2 Proksch E, Brandner JM, Jensen JM. The skin: An indispensable barrier. Exp Dermatol. 2008; doi: 10.1111/j.1600-0625.2008.00786.x.

3 Madison KC. Barrier function of the skin: “la raison d’être” of the epidermis’’. J Invest Dermatol. 2003; doi: 10.1046/j.1523-1747.2003.12359.x.

4 Wood LC, Jackson, SM, Elias PM, et al. Cutaneous barrier perturbation stimulates cytokine production in the epidermis of mice. J Clin Invest. 1992; doi: 10.1172/JCI115884.

5 Wang XP, Schunck M, Kallen KJ, et al. The interleukin-6 cytokine system regulates epidermal permeability barrier homeostasis. J Invest Dermatol. 2004; doi: 10.1111/j.0022-202X.2004.22736.x

6 Jensen JM, Schütze S, Förl M, et al. Roles for tumor necrosis factor receptor p55 and sphingomyelinase in repairing the cutaneous permeability barrier. J Clin Invest. 1999; doi: 10.1172/JCI5307.

7 Man MQ, Choi EH, Schmuth M, et al. Basis for improved permeability barrier homeostasis induced by PPAR and LXR activators: Liposensors stimulate lipid Synthesis, lamellar body secretion, and post-secretory lipid processing. J Invest Dermatol. 2006; 126: doi: 10.1038/sj.jid.5700046

8 Mizutani Y, Sun H, Ohno Y, et al. Cooperative synthesis of ultra long-chain fatty acid and ceramide during keratinocyte differentiation. PLOS ONE. 2013; doi: 10.1371/journal.pone.0067317.

9 Hong JW, Lee WJ, Hahn SB, et al. The effect of human placenta extract in a wound healing model. Ann Plast Surg. 2010; doi: 10.1097/SAP.0b013e3181b0bb67.

10 Hong KB, Park Y, Kim JH, et al. Effects of porcine placenta extract ingestion on ultraviolet B-induced skin damage in hairless mice. Korean J Food Sci Anim Resour. 2015; doi: 10.5851/kosfa.2015.35.3.413.

11 O’Keefe EJ, Payne RE, Russell N. Keratinocyte growth-promoting activity from human placenta. J Cell Physiol. 1985; doi: 10.1002/jcp.1041240312.

12 Cho HR, Ryou JH, Lee JW, et al. The effects of placental extract on fibroblast proliferation. J Cosmet Sci. 2008; 59: 195–202

13 Aioi A, Muromoto R, Mogami S, et al. Porcine placenta extract reduced wrinkle formation by potentiating epidermal hydration. J Cosmet Dermatol Sci Appl. 2021; doi: 10.4236/jcdsa.2021.112011

14 Sassa T, Hirayama T, Kihara A. Enzyme activities of the ceramide synthases CERS2– 6 are regulated by phosphorylation in the C-terminal region. J Biol Chem. 2016; doi: 10.1074/jbc.M115.695858.

15 Mullen TD, Hannun YA, Obeid LM. Ceramide synthases at the centre of sphingolipid metabolism and biology. Biochem J. 2012; doi: 10.1042/BJ20111626.

16 Levy M, Futerman AH. Mammalian ceramide synthases. IUBMB Life. 2010; doi: 10.1002/iub.319.

17 Nicholson KM, Anderson NG. The protein kinase B/Akt signaling pathway in human malignancy. Cell Signal. 2002; doi: 10.1016/s0898-6568(01)00271-6.

18 Persad S, Attwell S, Gray V, et al. Regulation of protein kinase B/Akt-serine 473 phosphorylation by integrin-linked kinase: Critical roles for kinase activity and amino acids arginine 211 and serine 343. J Biol Chem. 2001; doi: 10.1074/jbc.M102940200.

19 Di-Poï N, Tan NS, et al. Antiapoptotic role of PPARβ in keratinocytes via transcriptional control of the Akt1 signaling pathway. Mol Cell. 2002; doi: 10.1016/s1097-2765(02)00646-9.

20 Dibble CC, Cantley LC. Regulation of mTORC1 by PI3K signaling. Trends Cell Biol. 2015; doi: 10.1016/j.tcb.2015.06.002.

21 Buerger C, Shirsath N, Lang V, et al. Inflammation dependent mTORC1 signaling interferes with the switch from keratinocyte proliferation to differentiation. PLOS ONE. 2017; doi: 10.1371/journal.pone.0180853.

22 Bromberg, J, Darnell, JE Jr. The role of STATs in transcriptional control and their impact on cellular function. Oncogene. 2000; doi: 10.1038/sj.onc.1203476.

23 Levy DE, Lee CK. What does Stat3 do? J Clin Invest. 2002; doi: 10.1172/JCI15650.

24 Sano S, Itami S, Takeda K, et al. Keratinocyte-specific ablation of Stat3 exhibits impaired skin remodeling, but does not affect skin morphogenesis. EMBO J. 1999; doi: 10.1093/emboj/18.17.4657.

25 Zhou J, Wulfkuhle J, Zhang H, et al. Activation of the PTEN/mTOR/STAT3 pathway in breast cancer stem-like cells is required for viability and maintenance. Proc Natl Acad Sci U S A. 2007; doi: 10.1073/pnas.0702596104.

